# Imatinib Reduces Right Ventricular Systolic Pressure Independent of Arterial or Venous Remodeling in an Inflammatory Murine Model of Pulmonary Hypertension

**DOI:** 10.64898/2026.05.05.723006

**Authors:** Madeleine McGlynn, Lea C. Steffes, Anvi Shah, Joceline Morales, Maya E. Kumar

## Abstract

Pulmonary arterial hypertension is a progressive, fatal disease driven by pathologic vascular remodeling including arterial medial hypertrophy, occlusive neointimal lesion formation, and venous muscularization. Current vasodilatory therapies improve hemodynamics but do not reverse established remodeling. Imatinib mesylate, a tyrosine kinase inhibitor targeting the PDGF-PDGFR signaling axis, has been proposed as an anti-remodeling therapy for pulmonary arterial hypertension and has demonstrated hemodynamic benefit in both preclinical models and clinical trials. However, prior preclinical models lack the neointimal lesions characteristic of human disease, effects on venous remodeling have not been examined, and direct histologic assessment in human trials is precluded by the invasiveness of serial lung biopsy. Here, leveraging the house dust mite mouse model of pulmonary hypertension, which recapitulates medial thickening, neointimal lesion formation, and venous muscularization, we rigorously evaluate the anti-remodeling and hemodynamic effects of imatinib during two defined remodeling stages: neointimal lesion growth and neointimal lesion maintenance.

Imatinib treatment significantly reduced right ventricular systolic pressure at both stages. Despite this hemodynamic improvement, quantitative vessel-level analysis of over 1,700 arteries and 1,200 veins revealed no significant effect of imatinib on arterial medial thickness, neointimal lesion growth, neointimal lesion maintenance, or venous muscularization across any vessel size class. These findings dissociate imatinib’s hemodynamic benefit from structural vascular remodeling and suggest that imatinib functions primarily as a pulmonary vasodilator rather than an anti-remodeling agent.

## Introduction

Pulmonary arterial hypertension (PAH) is a progressive and fatal disease of the pulmonary vascular bed, with median 5-year survival around 60% despite current therapeutic options(1). PAH pathology is driven by ongoing vascular remodeling causing small vessel obliteration. This remodeling includes arterial medial hypertrophy and the formation of occlusive neointimal lesions that progressively obstruct the vessel lumen(2). Venous remodeling is increasingly recognized across multiple forms of pulmonary hypertension, including PAH(2). These changes result in sustained elevated pulmonary arterial pressures, ultimately leading to right ventricular failure and even death. The majority of PAH therapies are vasodilators targeting the prostacyclin, endothelin, or nitric oxide pathways. While these agents improve functional capacity and hemodynamics, they neither slow disease progression nor reverse established vascular remodeling. True anti-remodeling therapies that directly target the cellular and molecular mechanisms driving pathologic vascular remodeling are needed to decrease morbidity and mortality in this population(3–5).

Both medial hypertrophy and neointimal lesion formation are processes driven by proliferation of smooth muscle lineages(6–9). For this reason, platelet-derived growth factor (PDGF) signaling has been a compelling anti-remodeling target in PAH since PDGF signaling promotes vascular smooth muscle cell (VSMC) proliferation and migration(10). Notably, genetic ablation of PDGF signaling blunts arterial muscularization and RVSP elevations in the mouse hypoxia model of pulmonary hypertension(11). Imatinib mesylate (Gleevec, STI-571), a tyrosine kinase inhibitor originally developed as a BCR-Abl inhibitor, is a potent inhibitor of the PDGF-PDGFR signaling axis(12). Preclinical work by Schermuly and colleagues demonstrated that imatinib reduced right ventricular systolic pressure (RVSP) and thinned established arterial muscularization in the monocrotaline rat and hypoxia mouse models(13), findings subsequently replicated by other studies(14–16). Importantly, however, these models do not recapitulate the complex occlusive neointimal lesions of human disease, and venous remodeling was not interrogated.

A subsequent randomized controlled trial of imatinib in patients with advanced PAH reported promising reductions in pulmonary vascular resistance but was discontinued due to serious adverse events including subdural hematomas(17). More recent efforts using lower dose oral regimens, inhaled formulations and other tyrosine kinase inhibitors with more specific binding patterns have yielded equivocal results, leaving the clinical utility and true anti-remodeling capacity of imatinib unresolved(18–22). No histologic assessment of vascular remodeling in human lung tissue before and after imatinib treatment has been reported. Nevertheless, tyrosine kinase inhibitors remain among the most actively investigated therapeutic targets in PAH, highlighting the need for a comprehensive understanding of imatinib’s effects on pulmonary vascular remodeling(23).

The house dust mite (HDM) mouse model of pulmonary hypertension recapitulates medial thickening, neointimal lesion formation and venous muscularization characteristic of human PAH(6), providing a robust platform for evaluating a drug’s effect on both pre-and post-capillary remodeling. Further, remodeling in this system progresses through precisely defined stages, including medial thickening, neointimal establishment, neointimal lesion growth and maintenance of existing lesions. By staggering treatment such that animals are exposed to drug during discrete remodeling windows, we can assess a drug’s effect on each distinct stage in vessel remodeling (Figure 1A)(6, 24). Critically, vessel-by-vessel quantification of remodeling parameters across individual animals enables rigorous analysis of a drug’s effects, distinguishing responses in the arterial medial, neointimal, and venous compartments rather than relying on the global measures of muscularization or non-, partial-and fully-muscularized vessel assessments often used in preclinical studies.

**Figure 1:**
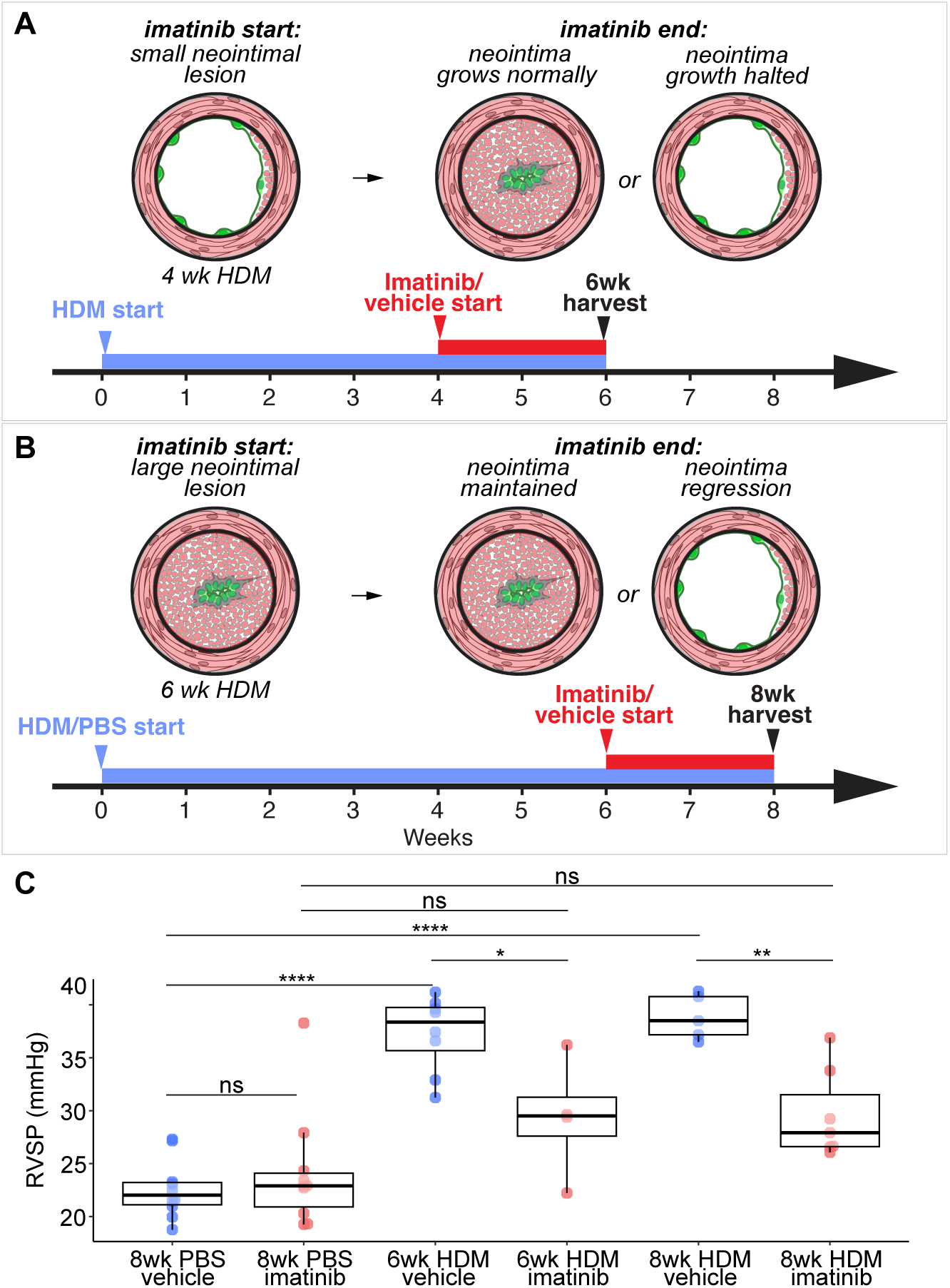
Study design and significant RVSP improvement in response to 2 weeks of imatinib treatment in the HDM mouse model. Schematics illustrating study design with arms to test both prevention of neointimal growth by imatinib, **A**, and regression of established neointimal lesions in response to imatinib treatment, **B**. HDM is given for either 6 weeks or 8 weeks (blue bars), where in the final 2 weeks animals are also exposed to either imatinib or vehicle (red bars). Following 4 weeks of HDM exposure (“imatinib start” in **A**) pulmonary arteries have small neointimal lesions that grow over the next two weeks to fully occlude the lumen (“imatinib start” in **B**) after which they will maintain their size if HDM exposure continues. Potential outcomes of the study (“imatinib end”) are schematized. **C**, RVSP is significantly elevated following both 6 and 8 weeks of HDM exposure. HDM exposed animals treated with imatinib show significant improvement in RVSP compared with vehicle treated animals, consistent with previous reports. Colored dots are measurements from individual animals, overlaid box plots indicate quartiles and medians for each group. Differences in RVSP across groups were assessed by one-way ANOVA followed by Tukey honest significant difference post-hoc test for pairwise comparisons. Normality was assessed within each group using the Shapiro-Wilk test and homogeneity of variance was confirmed using Levene’s test. One group (8-week PBS imatinib) did not meet the normality assumption (Shapiro-Wilk p = 0.006); all other groups passed. Schematics of artery cross sections in **A**&**B** show endothelium in green, internal and external elastic laminae with heavy black lines, and media and neointima in red. Media is located between internal and external elastic laminae; neointima is located between the internal elastic lamina and the basal edge of the endothelium. Significance thresholds: ns = not significant (p ≥ 0.05); * p < 0.05; ** p < 0.01; **** p < 0.0001.

In this study, we leveraged the HDM model of PH to test the effect of imatinib treatment on pulmonary vascular remodeling, including medial thickening, neointimal growth, and neointimal maintenance in arteries, and muscularization in veins with vessel-level resolution. While imatinib significantly reduced RVSP, we observed no significant change in any vascular remodeling parameter across any vessel type, vascular compartment or diameter class. Timed drug treatment showed that imatinib treatment neither slowed neointimal growth during lesion development, nor induced neointimal regression in established disease. Our findings demonstrate that hemodynamic improvement in response to imatinib treatment is not due to an anti-remodeling effect in this model and instead suggests that imatinib functions as a pulmonary vasodilator.

## Methods

### Animals and HDM Model

10-week-old female BALB/c mice were obtained from Charles River (strain 028). For the HDM model of PH, lyophilized HDM extract (Stallergenes Greer, XPB70D3A25) was diluted to a concentration of 20 μg Derp1 protein per 50 μl in sterile PBS, and aliquots were stored at −20°C. Once thawed, an aliquot was not refrozen, though excess solution could be held overnight at 4°C for use the following day. Fifty microliters of HDM solution were delivered intranasally to animals briefly anesthetized by isoflurane inhalation (2.5% isoflurane nebulized in 2L 100% oxygen per minute). Animals were dosed five days per week for the indicated duration. Control animals received an equal volume of sterile PBS delivered by the same route and following the same dosing schedule. Animals were harvested at 6 or 8 weeks of HDM exposure, corresponding to the periods of active neointimal expansion and established neointimal maintenance, respectively (Figure 1A). All animal experiments were performed under protocols approved by the Stanford University Institutional Animal Care and Use Committee.

### Imatinib Treatment

Imatinib methanesulfonate salt (LC Laboratories, Cat. No. I-5508) was dissolved in sterile distilled water at a concentration of 20 mg/mL and stored at 4 degrees Celsius for up to 5 days. Animals received imatinib at 100 mg/kg/day by oral gavage. Vehicle-treated animals received sterile distilled water delivered identically. Treatment was administered daily for two weeks during one of two defined windows: weeks 5 and 6 of HDM exposure, corresponding to the period of active neointimal expansion, or weeks 7 and 8 of HDM exposure, corresponding to the period of neointimal maintenance. Animals were weighed daily throughout the treatment period. Animals were excluded from analysis if body weight dropped more than 20% from baseline during the dosing window, though no animals met this exclusion criterion.

### Hemodynamic Assessment

RVSP was measured by closed chest right jugular vein catheterization performed under isoflurane anesthesia (induction conditions: 2.5% isoflurane nebulized in 2L 100% oxygen per minute in an induction box; maintenance conditions: 2.5% isoflurane nebulized in 400 ml 100% oxygen per minute to an active scavenging nose cone system). All catheterizations were performed by blinded lab members trained in this procedure. All animal experiments were performed under protocols approved by the Stanford University Institutional Animal Care and Use Committee.

### Tissue Collection, Cryopreservation and Sectioning

Animals were euthanized following RVSP measurement by severing the descending aorta while under isoflurane anesthesia. Blood was washed from the pulmonary circulation by slow (<100 μl per 10 seconds) perfusion of sterile phosphate buffered saline (PBS; Thermo-Fisher, J61196.AP) into the right ventricle. Airways were inflated with 2% low melting point agarose (Invitrogen, 16520) prepared in sterile PBS, and heart-lung blocks were removed from the chest cavity. Tissue for antibody staining was submerged in 40 ml ice-cold 4% paraformaldehyde solution in PBS (EMS, 15710) and allowed to fix for 4 hours at 4°C with gentle shaking. Fixed lobes were cryopreserved in 30% sucrose in PBS overnight, embedded in OCT compound (Sakura, 4583), frozen at −80°C and 20 μm sections were collected using a Leica CM3050S cryostat. Sections were stored at −80°C prior to staining.

### Immunohistochemistry

Slide labels were obscured and replaced with random three-letter codes prior to staining, blinding sample identity for the duration of staining, imaging, and quantification. Cryosections were stained following standard protocols as previously described(6). Briefly, slides were thawed, washed in PBS with 0.1% Tween-20, and blocked for at least 30 minutes in a preblock solution consisting of 0.3% Triton X-100, 5% serum, and 1.5% BSA. Slides were then incubated overnight at room temperature in primary antibody solution diluted in preblock, washed the following day, and incubated for 45 minutes in secondary antibody solution. Nuclei were stained with DAPI (1:1000; Invitrogen, D1306) and slides were mounted with Prolong Gold Antifade (Invitrogen, P36930). Elastin layers were visualized using fluorescent hydrazide dye; stock solutions were prepared by dissolving 1 mg hydrazide-A633 (Invitrogen, A30634) in 2 ml diH2O, used at 1:500 in the primary antibody mix. Primary antibodies used: mouse anti-SMA-Cy3 (Sigma, C6198; 1:200), rat anti-mouse CD31 (BD Pharmingen, 553370; 1:500).

Full lobe high-resolution four-color tiled confocal images were acquired with a 25x objective on a Zeiss 880 Examiner confocal microscope and minimally processed using Zen Black software (Carl Zeiss AG).

### Quantitative Image Analysis

All image analysis was performed under blinded conditions. For each animal, arterial cross sections between 25 to 150 μm in average external diameter and veins greater than 25 μm in average external diameter were quantitated from at least three tiled scans of full lobe sections, collecting a range of 41-89 artery and 21-84 vein measurements per animal (Figure S1). Arteries were distinguished from veins by the presence of both an internal and external elastic lamina. For each artery, two orthogonal measurement axes were drawn, and the following parameters were recorded in μm: external vessel diameter, medial thickness, and neointimal thickness when present (Figure S1). Medial thickness was defined as the ACTA2+ cell layers spanning the distance from the inner margin of the external elastic lamina to the outer margin of the internal elastic lamina. Neointimal thickness was defined as the ACTA2+ cell layers from the inner margin of the internal elastic lamina to the basal edge of the endothelium. For each vein, two orthogonal measurement axes were drawn and the following parameters recorded: external vessel diameter and muscular thickness. Muscular thickness was defined as the ACTA2+ cell layers extending from the inner margin of the external elastic lamina to the basal edge of the endothelium. All measurements were collected manually with the assistance of Zen Black software (Carl Zeiss AG; Figure S1). Medial thickness is reported as an average of all four measurements per vessel. Neointimal lesion size is reported as an average of the four neointimal measurements per vessel and expressed as a percent of that vessel’s average external diameter.

Venous muscularization is reported as the average of the four muscle thickness measurements and expressed as a percent of that vessel’s average external diameter.

## Statistical Analysis

All analyses were performed in R using the outliers, car, lme4, lmerTest, and emmeans packages as detailed below. Remodeling parameters (medial thickness, percent neointima, and percent venous muscularization) were analyzed using linear mixed effects models to account for measurements of multiple vessels within individual animals. Estimated marginal means (emmeans) were extracted from each model using the emmeans package. All pairwise comparisons across the six experimental groups were computed for each outcome with Tukey honest significant difference adjustment. Comparisons of primary interest – vehicle versus imatinib within each exposure group (8-week PBS, 6-week HDM, and 8-week HDM) – were extracted from the full set of Tukey-adjusted pairwise comparisons and are reported in the main text; all 15 pairwise comparisons for each outcome are reported in Table S1.

Remodeling parameters are presented as emmeans in the text. Differences in RVSP across groups were assessed by one-way ANOVA followed by Tukey honest significant difference post-hoc test for pairwise comparisons. Normality was assessed within each group using the Shapiro-Wilk test and homogeneity of variance was confirmed using Levene’s test. One group (8-week PBS imatinib) did not meet the normality assumption (Shapiro-Wilk p = 0.006); all other groups passed. RVSP data reported in the text are presented as mean ± standard deviation. To assess for the presence of a high-responding animal subpopulation, per-animal mean medial thickness and percent neointima were calculated by averaging vessel-level measurements within each animal. Grubbs’ test was applied to per-animal means within each group and outcome to formally test for statistical outliers, with p > 0.05 taken as evidence against the presence of an outlier animal.

## Results

### Imatinib Treatment Reduces RVSP in the HDM Model

Preclinical and clinical trials have demonstrated significantly improved hemodynamics in PAH with imatinib treatment(13, 15–17, 20, 21). To assess the hemodynamic effect of imatinib in the HDM model of PH, RVSP was measured by right heart catheterization. As previously established, in the HDM model medial thickening and venous remodeling are established after 2 weeks of HDM exposure.

Neointimal founder cells are widely observed after 4 weeks of HDM exposure and rapidly proliferate in the following 2 weeks to form large lesions after 6 weeks of HDM exposure. These lesions are maintained indefinitely if HDM exposure continues(6). In this study six exposure groups were tested: animals exposed to HDM for 6 weeks that were given either imatinib or vehicle during weeks 5 and 6 to test for prevention of neointimal growth (Figure 1A); animals exposed to HDM for 8 weeks that were given either imatinib or vehicle during weeks 7 and 8 to test for reversal of neointimal lesions (Figure 1B); and control animals exposed to PBS for 8 weeks that were given either imatinib or vehicle during weeks 7 and 8 (Figure 1B). Reversal of arterial medial thickening and venous muscularization by imatinib is tested by both HDM durations.

RVSP measurements were collected in all groups at harvest (Figure 1C). PBS controls exposed to vehicle had significantly lower RVSP than both vehicle exposed HDM groups (22.6 ± 2.8 mmHg vs 37.3 ± 3.6 mmHg for 6-week HDM vehicle, p < 0.0001; and vs 38.8 ± 2.1 mmHg for 8-week HDM vehicle, p < 0.0001), confirming HDM-induced pulmonary hypertension. Imatinib treatment significantly reduced RVSP compared to vehicle controls in both HDM exposure groups (6wk: 29.4 ± 5.7 vs 37.3 ± 3.6 mmHg, p = 0.039; 8wk: 29.6 ± 4.2 vs 38.8 ± 2.1 mmHg, p = 0.006). No significant difference in RVSP was observed between PBS control animals receiving vehicle and PBS control animals receiving imatinib (p = 0.962). No significant difference was found between PBS imatinib treated mice and either HDM imatinib treated group (p = 0.299 vs 6wk HDM imatinib; p = 0.109 vs 8wk HDM imatinib; n = 10 vehicle, 10 imatinib for 8-week PBS; n = 8 vehicle, 4 imatinib for 6-week HDM; n = 5 vehicle, 7 imatinib for 8-week HDM). These data demonstrate a significant reduction in RVSP following imatinib treatment in the HDM model, consistent with prior preclinical reports and clinical trials.

### Imatinib does not reverse established medial thickness, prevent neointimal growth or reverse established neointimal lesions

To determine if and how imatinib affects medial thickness, neointimal growth, and neointimal maintenance, pulmonary arteries were quantified at the individual vessel level across all exposure groups and treatment windows. Samples were given a random three letter code to blind researchers, and cryosections were stained to identify endothelium, smooth muscle, elastic laminae and neointima and imaged by high resolution confocal scans of full lobe sections (Figure S1). Vessel parameters were collected by hand with the assistance of image analysis software by blinded lab personnel from 1,764 arteries in 35 animals. Arteries between 25 and 150 µm in average diameter were included in the analysis.

Vessel diameter did not differ significantly between vehicle and imatinib-treated animals in any exposure group (Figure S2). As expected, HDM exposure was associated with significantly increased medial thickness compared to PBS controls in vehicle-treated animals (Figure 2A&B emmeans: 8wk PBS vehicle 3.79µm vs 6wk HDM vehicle 6.78µm, p = 0.004; 8wk PBS vehicle 3.79µm vs 8wk HDM vehicle 7.92µm, p < 0.0001). However, medial thickness did not differ significantly between vehicle and imatinib-treated animals in any group (emmeans: 8-week PBS vehicle 3.79µm vs imatinib 2.67µm, p = 0.741; 6-week HDM vehicle 6.78µm vs imatinib 7.81µm, p = 0.822; 8-week HDM vehicle 7.92µm vs imatinib 7.62µm, p = 0.998).

**Figure 2:**
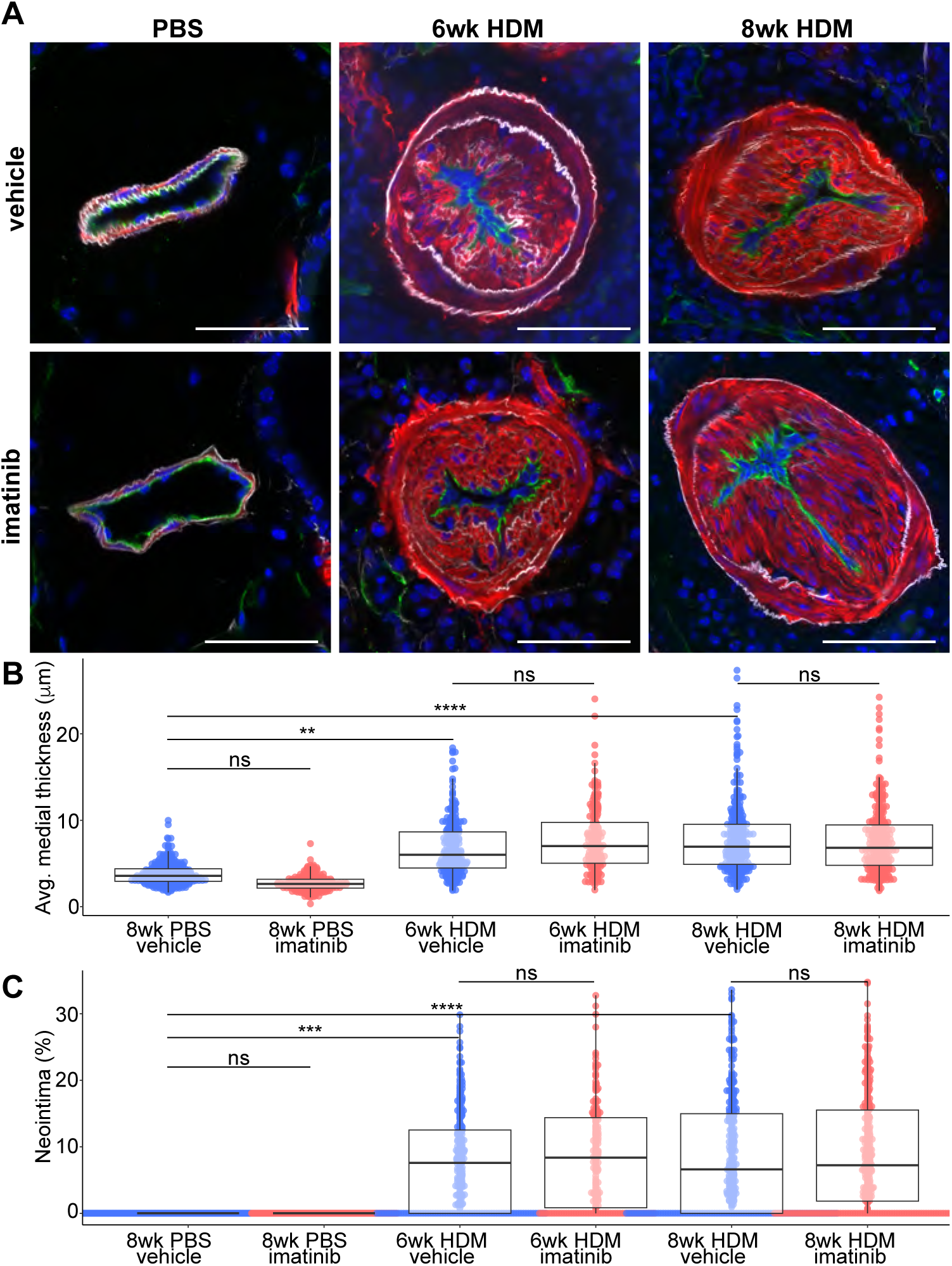
Compartment specific quantitation of artery remodeling shows no thinning of the media, slowing of neointimal growth, or neointimal lesion regression in response to imatinib treatment. **A**, Confocal images of arteries stained to highlight endothelium (CD31, green), elastin (hydrazide dye, white), media and neointima (smooth muscle α-actin, red) and nuclei (DAPI, blue) from representative arteries of each treatment group are shown. Measurements collected from images of 1,764 arteries from 35 animals show that, **B**, the smooth muscle media thickens in response to HDM exposure but shows no reduction in medial thickness between imatinib treated and vehicle treated animals of either HDM exposure duration. **C**, Neointimal lesions are not present in PBS controls and are present following both 6 and 8 weeks of HDM, but no difference in neointimal size was detected between imatinib treated and vehicle treated animals at either HDM duration. Colored dots are the averaged measurements from a single artery, overlaid box plots indicate quartiles and medians for each group. Remodeling parameters (medial thickness and percent neointima) were analyzed using linear mixed effects models to account for measurements of multiple vessels within individual animals. Pairwise comparisons across the six experimental groups were computed for each outcome with Tukey honest significant difference adjustment. Scale bars are 50 µm. Significance thresholds: ns = not significant (p ≥ 0.05); ** p < 0.01; *** p < 0.001; **** p < 0.0001.

Neointimal lesions were absent in all PBS-exposed animals (Figure 2C). Among HDM-exposed animals, percent neointimal thickness did not differ significantly between vehicle and imatinib-treated animals at either timepoint (emmeans: 6-week HDM vehicle 8.08% vs imatinib 8.50%, p = 1.000; 8-week HDM vehicle 8.87% vs imatinib 9.22%, p = 1.000; n = 7 vehicle, 5 imatinib for 8-week PBS; n = 6 vehicle, 4 imatinib for 6-week HDM; n = 7 vehicle, 7 imatinib for 8-week HDM).

Taken together, imatinib treatment did not alter arterial medial thickness or percent neointimal thickness either during lesion growth or lesion maintenance, dissociating the significant hemodynamic effect of imatinib treatment from any measurable change in structural arterial vascular remodeling.

### Imatinib does not reduce pulmonary arterial remodeling across vessel size subpopulations

To determine whether imatinib’s effect on arterial remodeling might be restricted to a specific vessel size range, medial thickness and percent neointimal thickness were re-examined within three artery size ranges: 25–50 µm, 51–100 µm, and 101–150 µm in average diameter (Figure 3A). No significant difference in medial thickness was detected between vehicle and imatinib-treated animals in any size range at either HDM timepoint (Figure 3B; 25–50 µm: 6wk HDM p = 0.988, 8wk HDM p = 0.319; 51–100 µm: 6wk HDM p = 0.143, 8wk HDM p = 1.000; 101–150 µm: 6wk HDM p = 0.992, 8wk HDM p = 0.999). Similarly, percent neointimal thickness did not differ significantly between vehicle and imatinib-treated animals in any size range at either timepoint (Figure 3C; 25–50 µm: 6wk HDM p = 0.733, 8wk HDM p = 0.602; 51–100 µm: 6wk HDM p = 0.979, 8wk HDM p = 1.000; 101–150 µm: 6wk HDM p = 0.881, 8wk HDM p = 0.997). These findings confirm that the absence of a detectable imatinib effect on arterial remodeling when vessels of all sizes are pooled (Figure 2B&C) was not obscuring statistically significant effects in distinct vessel size subsets and demonstrates that structural arterial remodeling was unaffected by imatinib treatment across the full range of arteries examined.

**Figure 3:**
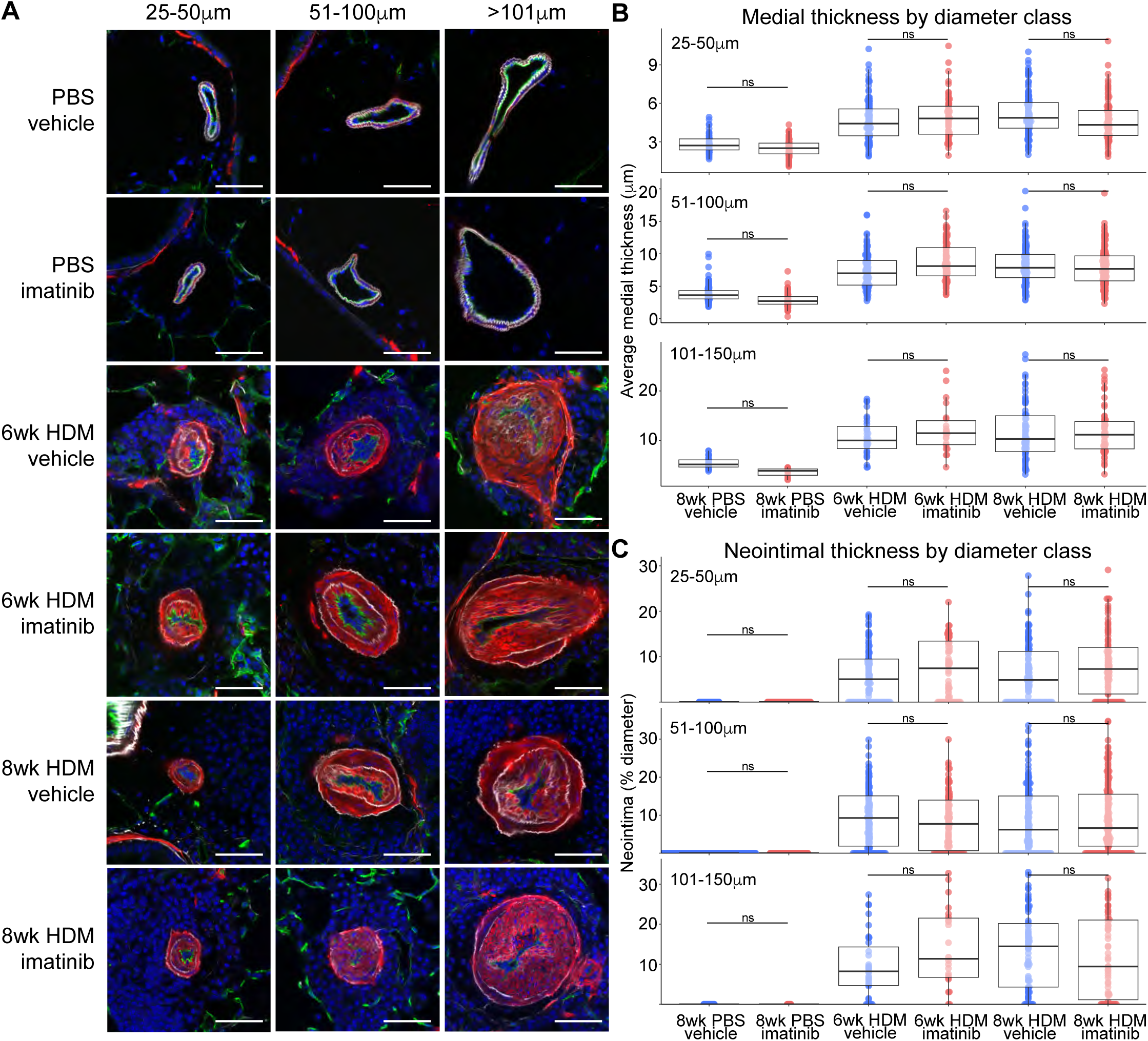
No effect of imatinib treatment on either media or neointima is revealed when arteries are subdivided by size. **A**, Confocal images of representative arteries from each size class (25-50 µm, 51-100 µm and 101-150 µm average external diameter) and treatment group stained to visualize endothelium (CD31, green), elastin (hydrazide dye, white), smooth muscle (smooth muscle α-actin, red), and nuclei (DAPI, blue) are shown. No changes in either medial thickness, **B**, or neointimal size, **C**, are detected in response to imatinib treatment in PBS controls or either HDM duration. Colored dots are the averaged measurements from a single artery, overlaid box plots indicate quartiles and medians for each group. Remodeling parameters (medial thickness and percent neointima) were analyzed using linear mixed effects models to account for measurements of multiple vessels within individual animals. All pairwise comparisons across the six experimental groups were computed for each outcome with Tukey honest significant difference adjustment. Scale bars are 50 µm. Significance thresholds: ns = not significant (p ≥ 0.05).

### Imatinib does not reverse established venous muscularization

Pulmonary venous remodeling is increasingly recognized across many forms of pulmonary hypertension, including those historically classified as predominantly precapillary(2). Advances in immunohistochemistry coupled with high-resolution confocal microscopy now permit reliable discrimination of small arteries and veins in tissue sections, making it possible to quantify this vascular compartment independently(6). Given that an anti-remodeling effect of imatinib confined to the venous circulation could be missed by analyses restricted to arteries alone, venous muscularization was quantified at the individual vessel level in each exposure group and treatment window, capturing 1,256 veins from 32 animals. Since venous muscularization begins within the first two weeks of HDM exposure and accumulates progressively thereafter(6), vein muscularization was present in both the 6-week and 8-week HDM groups at the time of imatinib administration, allowing assessment of imatinib’s effect on reversal of existing venous smooth muscle accumulation (Figure 4A).

**Figure 4:**
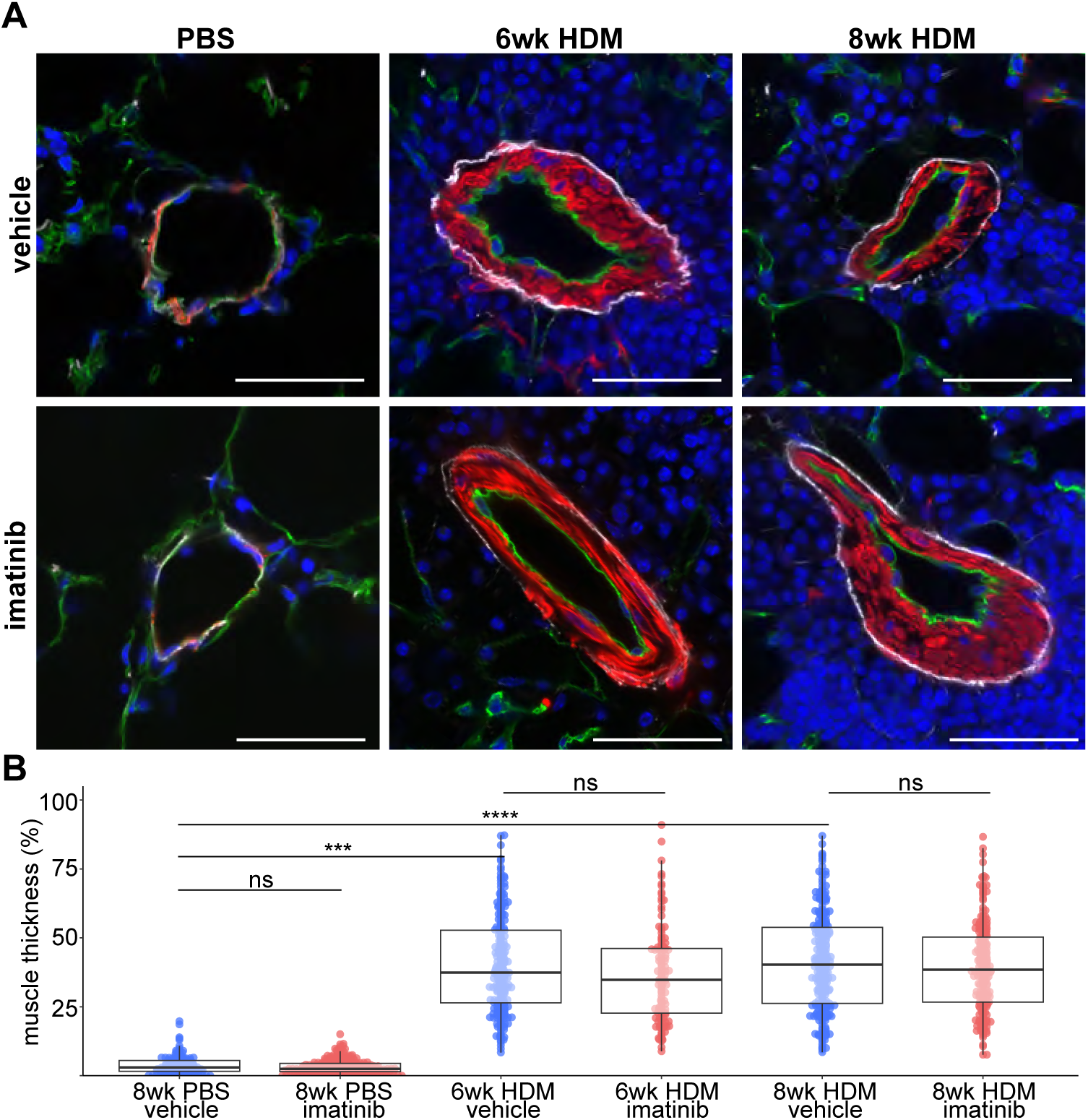
Venous muscularization is not reduced in response to imatinib treatment. **A**, Confocal images of representative veins from each treatment group stained to visualize endothelium (CD31, green), elastin (hydrazide dye, white), smooth muscle (smooth muscle α-actin, red), and nuclei (DAPI, blue) are shown. **B**, Measurements collected from images of 1,256 veins from 32 animals show that muscle thickness in vein walls increases in response to HDM and is not reduced in response to imatinib treatment in either the 6-or 8-week HDM group. Colored dots are the averaged measurements from a single vein, overlaid box plots indicate quartiles and medians for each group. Percent venous muscularization was analyzed using linear mixed effects models to account for measurements of multiple vessels within individual animals. All pairwise comparisons across the six experimental groups were computed for each outcome with Tukey honest significant difference adjustment. Scale bars are 50 µm. Significance thresholds: ns = not significant (p ≥ 0.05); *** p < 0.001; **** p < 0.0001.

As expected, HDM exposure was associated with significantly increased percent muscularization in veins compared to PBS controls in vehicle-treated animals (8wk PBS vehicle vs 6wk HDM vehicle: p < 0.001; 8wk PBS vehicle vs 8wk HDM vehicle: p < 0.0001), confirming HDM-induced venous remodeling at both stages (Figure 4B). However, percent muscularization did not differ significantly between vehicle and imatinib-treated animals at either HDM timepoint (emmeans: 6wk HDM vehicle 40.82% vs imatinib 37.04%, p = 0.996; 8wk HDM vehicle 46.34% vs imatinib 40.14%, p = 0.915). These findings indicate that imatinib does not reverse established pulmonary venous muscularization.

### Individual Animal Analysis Shows No High-Responder Subpopulation

To evaluate animal to animal variance within treatment groups and identify a possible “high responding” subset that might be masked by pooling data within groups, remodeling measurements were examined at the individual animal level. Per-animal medial thickness and percent neointimal thickness were examined for all HDM-exposed animals (Figure S3). Grubbs’ test identified no statistically significant outlier animals in any group for any outcome (all p > 0.05), confirming that the distribution of remodeling responses was statistically homogeneous within each group. These results argue against the possibility that a subset of imatinib-responsive animals was obscured by non-responding animals when data were pooled at the group level.

## Discussion

In this study, through rigorous quantitation of individual vessels across more than 1,700 arteries and 1,200 veins, coupled with hemodynamic assessment, we demonstrate that imatinib significantly reduces RVSP in the HDM model of pulmonary hypertension without a significant effect on any measure of pulmonary vascular remodeling. We tested for imatinib-dependent effects on both arterial and venous beds, and found no reduction in artery medial thickness, neointimal lesion growth, neointimal lesion maintenance or venous muscularization. Searching for if a significant effect was masked by pooling data, we found no effect when analyzing distinct artery size classes, or individual animals. This is the first assessment of imatinib’s capacity to inhibit neointimal growth, reverse established neointimal lesions, or reverse venous muscularization. Taken together, these findings dissociate the hemodynamic effects of imatinib from any change in arterial or venous remodeling, despite the compelling mechanistic rationale linking PDGF signaling inhibition to pathologic vascular smooth muscle proliferation in the lung.

Aligned with previous preclinical reports(13, 15, 16), we find a significant hemodynamic benefit to imatinib treatment. We anticipated our results would recapitulate prior findings of regression of medial thickness, and our primary aim was to extend that analysis by examining imatinib’s specific effect on the neointima and venous remodeling. However, we were unable to recapitulate the medial thinning reported in previous studies. This discrepancy may in part reflect differences in models, as this is the first study using the HDM model of PH which generates advanced, humanlike neointimal lesions in addition to medial thickening and thus differs meaningfully from the monocrotaline rat and hypoxia mouse models used in prior work. Methodologic differences between this and prior analyses may also contribute. Here, we capitalized on advancements in fluorescent immunohistochemistry and employed a staining panel optimized to distinguish arteries from veins and resolve key vessel compartments (media vs neointima) and used high resolution confocal microscopy to enable rigorous and precise assessment of remodeling parameters in thousands of arteries and veins across the entire macrovascular bed. These advances allowed us to assess remodeling with a level of granularity not previously applied to preclinical imatinib-PH studies.

The robust hemodynamic response to imatinib treatment we observed in the absence of any detectable improvement in vessel remodeling leads us to query alternate mechanisms that could cause such a drop in RVSP. Two possible mechanisms include imatinib induced vascular shunting or acute pulmonary vasodilation. We see no histological indication of collateral vessel formation around pulmonary arterial occlusions with imatinib treatment, though it is possible small endothelial sprouts would be missed in the 2D analysis performed, and literature implicates PDGF signaling – rather than its inhibition – as a driver of angiogenesis(25–31). More compellingly, multiple studies have demonstrated imatinib’s action as a potent and acute vasodilator in pulmonary arteries in rats(32), mice(33), guinea pigs(34) and human tissue(33). Its vasodilatory activity has been characterized across multiple experimental conditions including isolated pulmonary arterial rings, precision cut lung slices from both humans and rodent models, and in anesthetized catheterized animals. Prior studies show imatinib’s vasodilatory effect is endothelium-independent with proposed mechanisms including K+ channel activation and Ca2+ desensitization in VSMCs(32). Indeed, the rapid onset of RVSP reduction reported in imatinib-treated monocrotaline rats may be more consistent with imatinib functioning as a pulmonary vasodilator than a true anti-remodeling agent(13). PDGF-AB and PDGF-BB have long been appreciated as direct vasoconstrictors via calcium mobilization and thromboxane generation(35, 36) and imatinib’s inhibition of PDGF signaling may therefore be integral to its vasodilatory effect. Given this evidence, we conclude that imatinib is likely improving hemodynamics in our system by acting as a vasodilator. This suggests that imatinib and other tyrosine kinase inhibitors may have utility as adjunct vasodilators in PAH whose mechanism of action is distinct from available therapies.

There are limitations to the current study. We use a model driven by inflammation rather than direct endothelial injury or hypoxia, though both human disease(37) and the previously employed animal models are also marked by perivascular inflammatory cell infiltration(38, 39). As outlined above, our vascular analysis is performed exclusively in 2D sections, which may miss newly formed vascular branches acting as shunts in response to imatinib. We tested only a single dose of imatinib, though our dosing matched previous reports(13). Prevention of early remodeling events, including initiation of medial thickening, establishment of neointimal founder cells, and initiation of venous muscularization, were not tested, as patients with PAH typically present with established disease. Additionally, the two-week treatment windows, though sufficient for a significant reduction in RVSP, may not be sufficient treatment time to detect long term imatinib-dependent lesion regression. Future studies could employ a wider range of dosing regimens coupled with early prevention and chronic treatment arms.

In conclusion, imatinib reduces RVSP in our model through mechanisms distinct from vascular remodeling, consistent with acute pulmonary vasodilation rather than the anticipated antiproliferative effect. This distinction does not preclude imatinib from serving a meaningful therapeutic role in PAH as an orthogonal pulmonary vasodilator, but it does shift the mechanistic interpretation of its benefit. These findings underscore the importance of pairing hemodynamic assessment with rigorous, compartment specific histologic evaluation in preclinical studies. As novel therapies targeting vascular remodeling move through the preclinical pipeline(40), the HDM model offers the resolution required to rigorously evaluate anti-remodeling effects and identify drugs capable of truly modifying the underlying pathology in PAH.

## Declarations

### Ethical approval

All animal experiments were performed under protocols approved by the Stanford University Institutional Animal Care and Use Committee (IACUC-31869 and IACUC-27626).

### Guarantor

Maya E. Kumar serves as the Guarantor for this manuscript and takes responsibility for the integrity of the work as a whole.

### Author Contributions

M.M. developed the experimental plan, performed all pharmacologic inhibition experiments, tissue handling and sectioning, immunohistochemistry, confocal microscopy, and vessel quantification, initial data interpretation, and provided critical manuscript edits. L.C.S. wrote the initial manuscript, generated figures, performed additional confocal microscopy and vessel quantification, completed statistical analysis, and mentored M.M. through initial experimental design. A.S. performed venous quantification and critically reviewed the manuscript. J.M. performed immunohistochemistry and critically reviewed the manuscript. M.E.K. supervised experimental design, wrote the initial manuscript, generated figures, completed microscopy and additional vessel quantitation and provided senior oversight of the project.

### Author Contributions Note

M.M. and L.C.S. contributed equally to this work. The order of co-first authors was determined based on L.C.S.’s primary role in data analysis, figure generation, and initial manuscript drafting, while M.M. led experimental design, pharmacologic inhibition experiments, tissue generation and handling, and vessel quantification.

## Acknowledgement

We thank David Cornfield and the members of the Center of Excellence in Pulmonary Biology at Stanford University for their guidance and expertise regarding smooth muscle cell contractility and the pulmonary vasodilatory effects of imatinib. We thank Edda Spiekerkoetter for helpful discussions and for generously providing access to right heart catheterization facilities and for her technical expertise in hemodynamic assessment.

---

**Figure S1:**
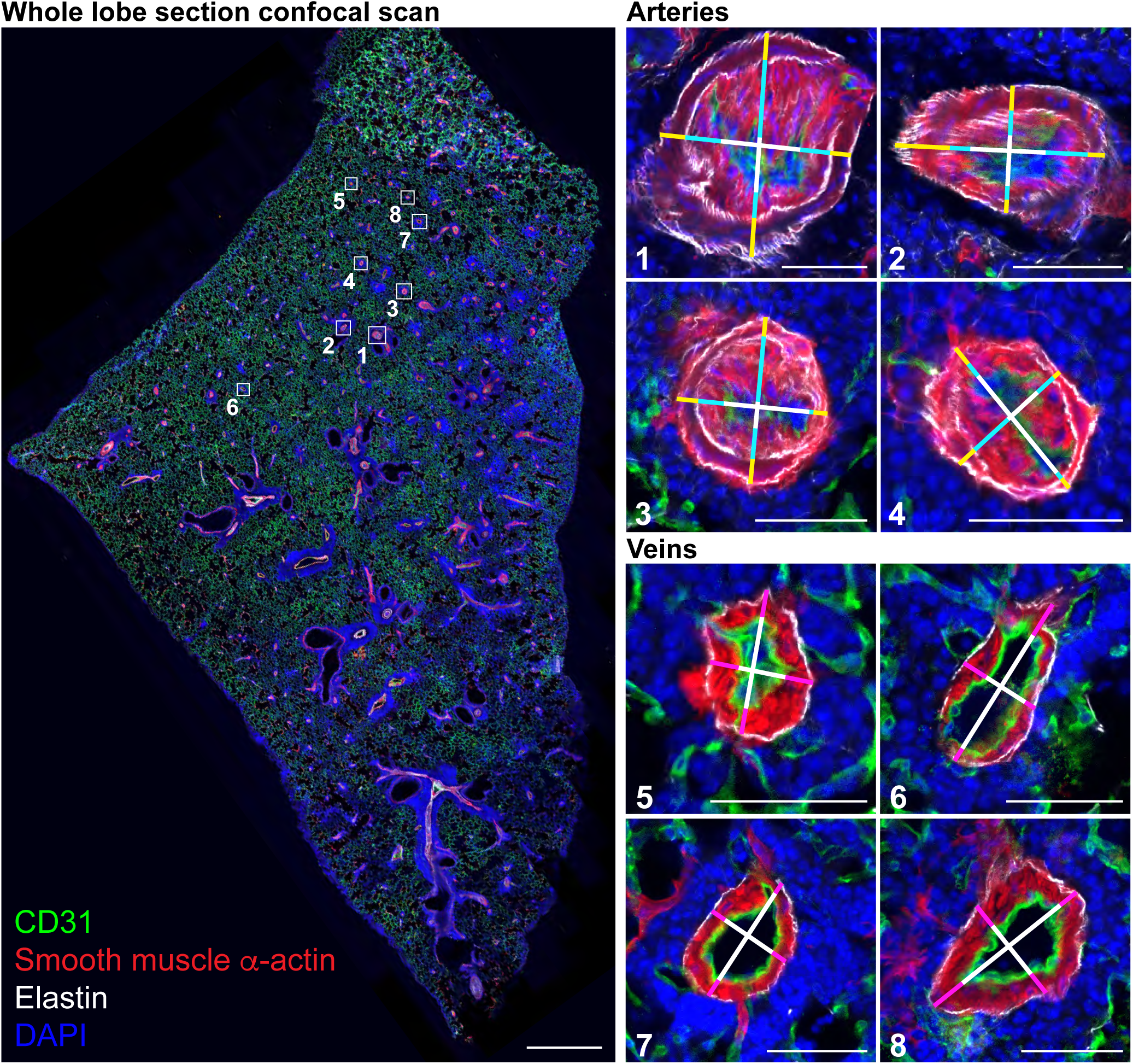
Unbiased image and data collection approach for quantitation of artery and vein remodeling parameters. Tiled high resolution confocal images were collected of whole lobe cryosections stained to highlight endothelium (CD31, green), elastin (hydrazide dye, white), smooth muscle α-actin (red), and nuclei (DAPI, blue), **left**. Blinded lab personnel scrolled through stitched whole section tiles in a grid pattern at full resolution, recording indicated vessel parameters along two orthogonal lines for all arteries between 25 and 150 µm in average diameter and all veins over 25 µm in average diameter. Selected arteries (**panels 1-4**) and veins (**panels 5-8**), indicated by the boxes at left, are highlighted at right. For arteries media (yellow line segment), neointima (cyan line segment), external diameter (full length of each line) were recorded. For veins muscle thickness (magenta line segment), and external diameter (full length of each line) were recorded. Scale bar, left, 1 mm. Scale bars, right, 50 µm.

**Figure S2:**
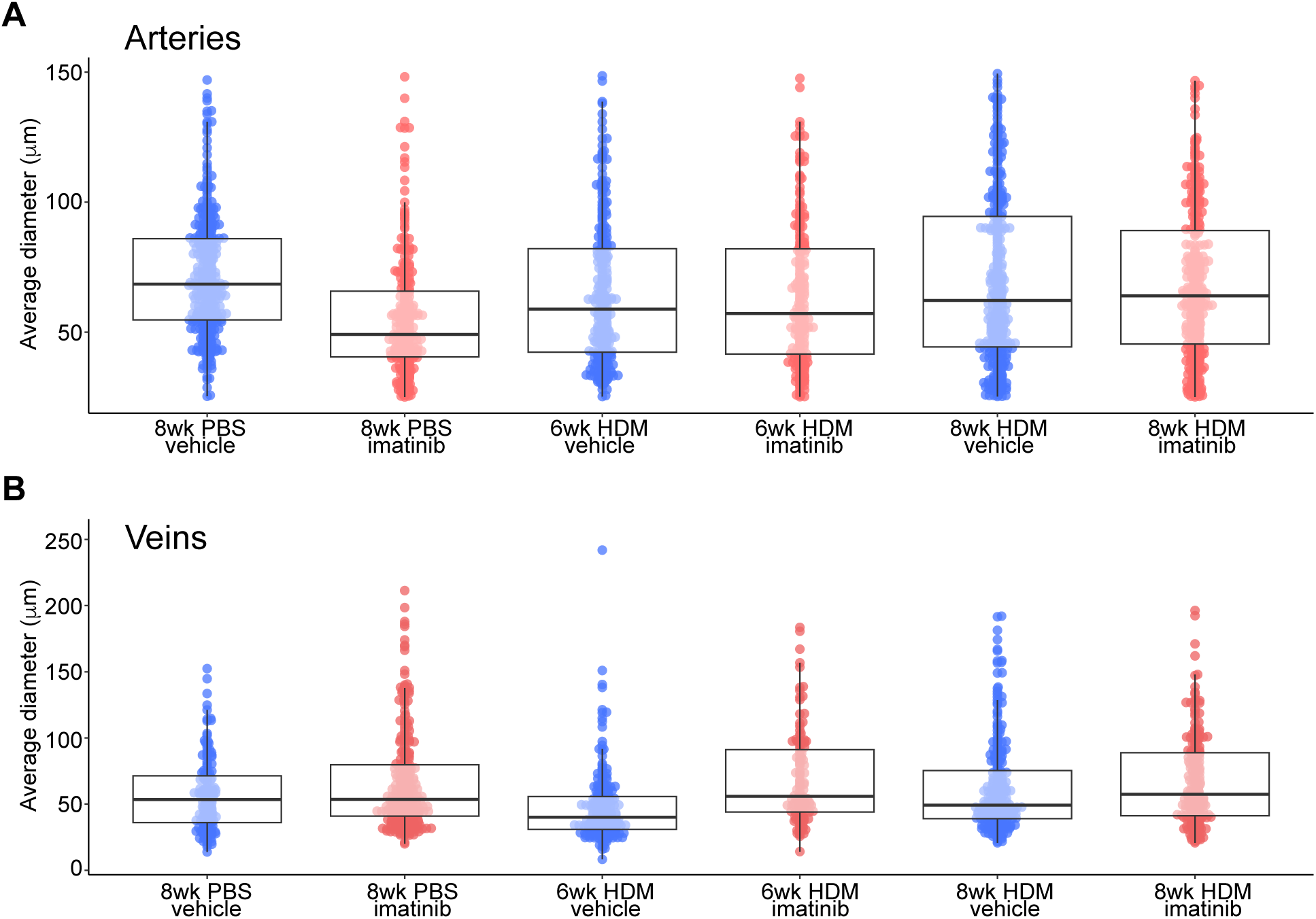
Average vessel diameter for each treatment group. No significant difference in average external diameter among either arteries, **A**, or veins, **B**, was detected among all sequence groups. Colored dots are the averaged measurements from a single vessel, overlaid box plots indicate quartiles and medians for each group. Pairwise comparisons across the six experimental groups were computed for each outcome with Tukey honest significant difference adjustment. No pairwise comparisons among arteries and among veins were significant (p ≥ 0.05).

**Figure S3:**
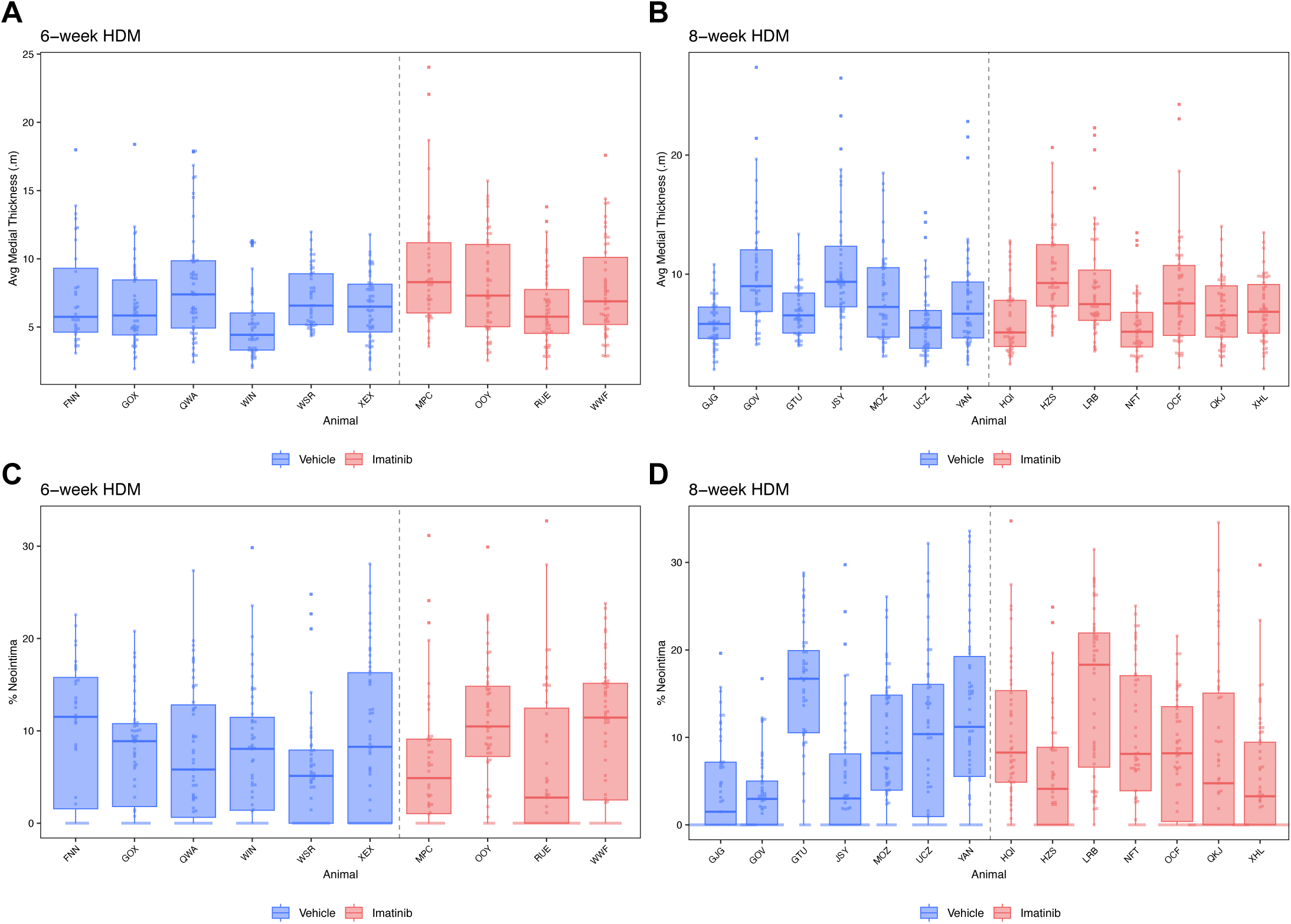
Analysis of arteries in individual animals found no imatinib ‘high responder’ subpopulation. No animals were found to be statistically significant outliers in either medial thickness (**A**&**B**), or neointimal lesion size (**C**&**D**). Colored dots are the averaged measurements from a single artery, overlaid box plots indicate quartiles and medians for each group. Per-animal mean medial thickness and percent neointima were calculated by averaging vessel-level measurements within each animal. Grubbs’ test was applied to per-animal means within each group and outcome to formally test for statistical outliers, with p > 0.05 taken as evidence against the presence of an outlier animal. Three letter codes indicate individual animals.

**Table S1:**
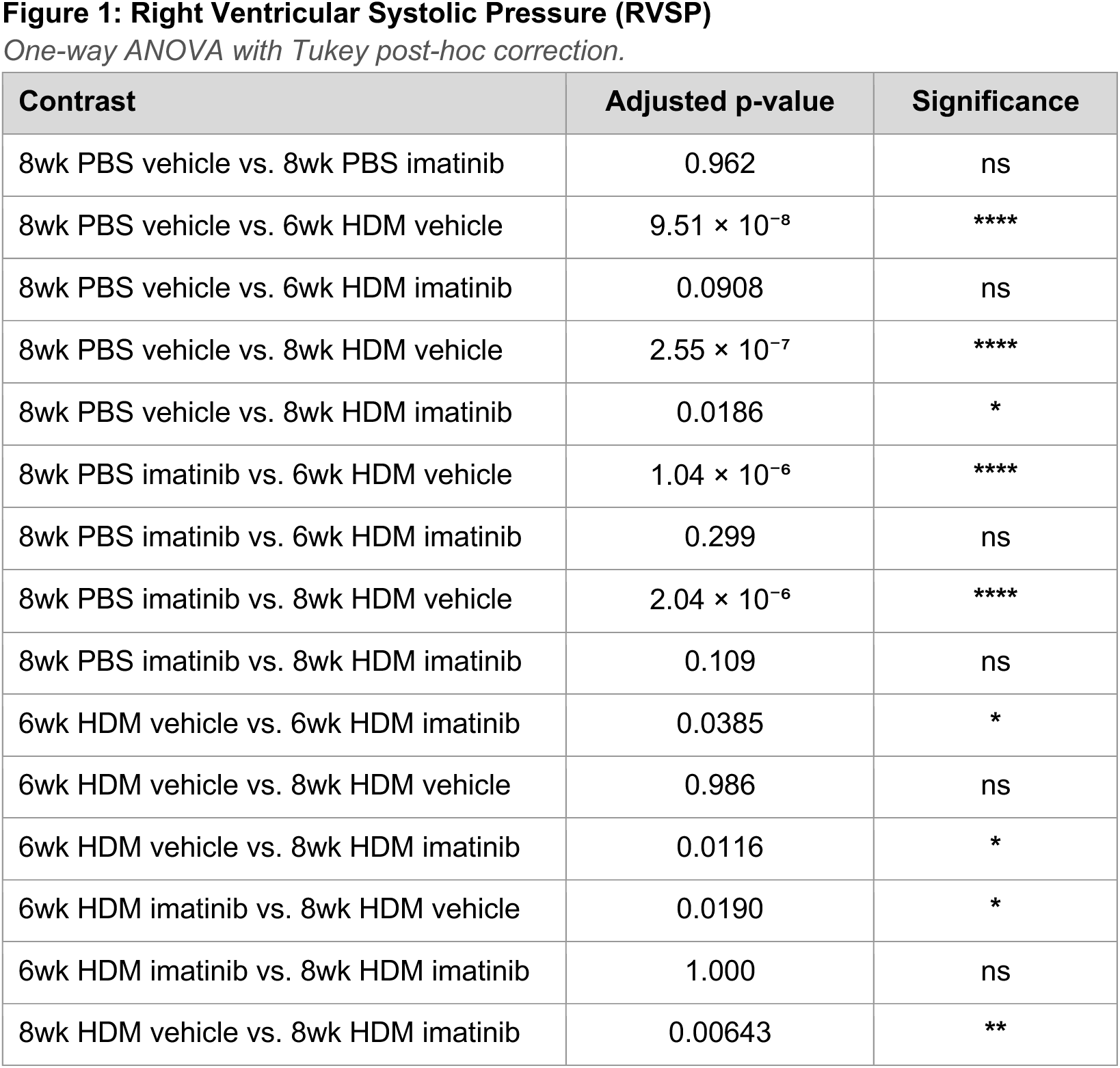
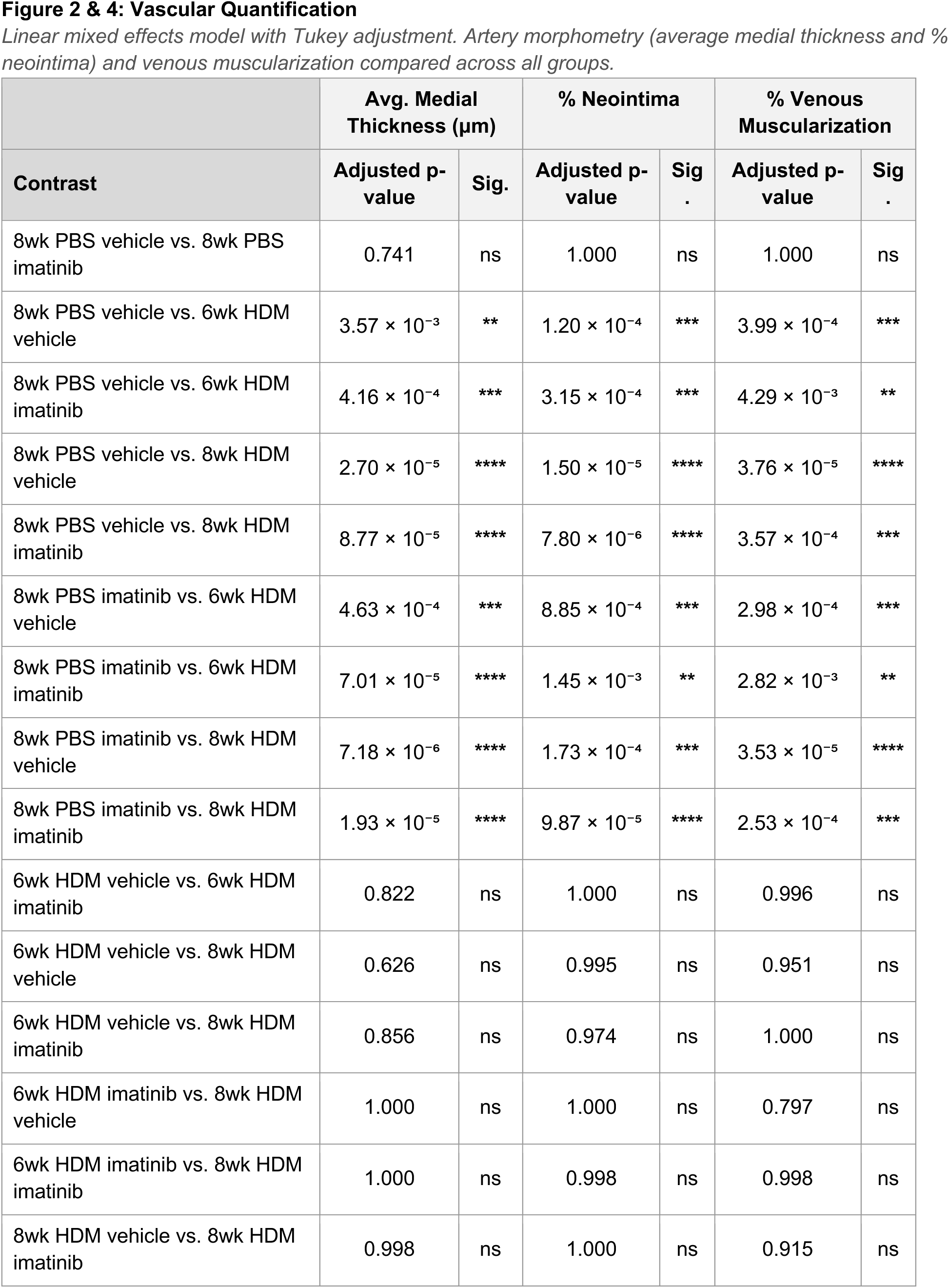
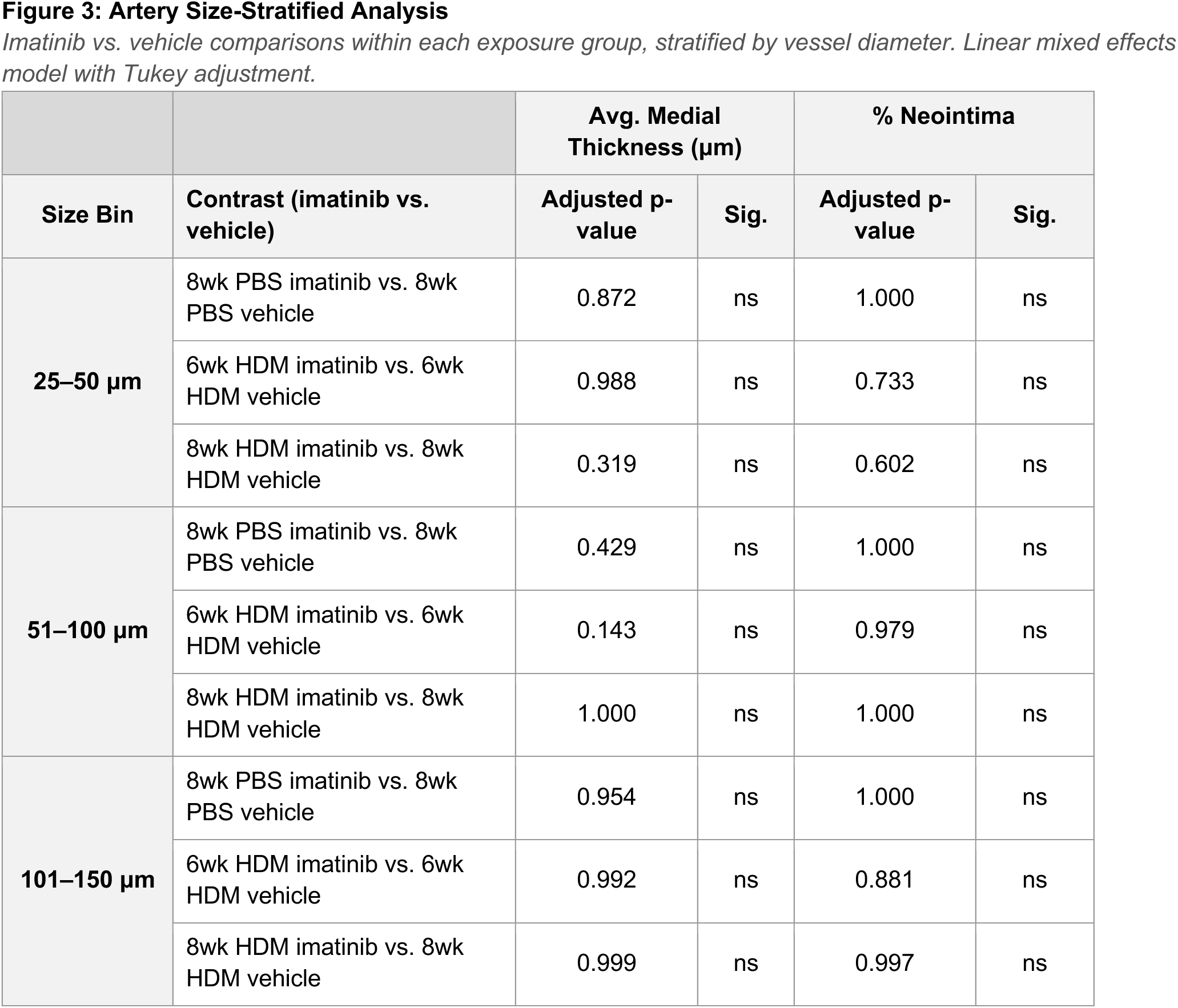
Statistical Comparisons for Figures 1-4. Significance thresholds: ns = not significant (p ≥ 0.05); * p < 0.05; ** p < 0.01; *** p < 0.001; **** p < 0.0001.

| Contrast | Adjusted p-value | Significance |
| --- | --- | --- |
| 8wk PBS vehicle vs. 8wk PBS imatinib | 0.962 | ns |
| 8wk PBS vehicle vs. 6wk HDM vehicle | $9.51 \times 10^{-8}$ | **** |
| 8wk PBS vehicle vs. 6wk HDM imatinib | 0.0908 | ns |
| 8wk PBS vehicle vs. 8wk HDM vehicle | $2.55 \times 10^{-7}$ | **** |
| 8wk PBS vehicle vs. 8wk HDM imatinib | 0.0186 | * |
| 8wk PBS imatinib vs. 6wk HDM vehicle | $1.04 \times 10^{-6}$ | **** |
| 8wk PBS imatinib vs. 6wk HDM imatinib | 0.299 | ns |
| 8wk PBS imatinib vs. 8wk HDM vehicle | $2.04 \times 10^{-6}$ | **** |
| 8wk PBS imatinib vs. 8wk HDM imatinib | 0.109 | ns |
| 6wk HDM vehicle vs. 6wk HDM imatinib | 0.0385 | * |
| 6wk HDM vehicle vs. 8wk HDM vehicle | 0.986 | ns |
| 6wk HDM vehicle vs. 8wk HDM imatinib | 0.0116 | * |
| 6wk HDM imatinib vs. 8wk HDM vehicle | 0.0190 | * |
| 6wk HDM imatinib vs. 8wk HDM imatinib | 1.000 | ns |
| 8wk HDM vehicle vs. 8wk HDM imatinib | 0.00643 | ** |

| Contrast | Avg. Medial Thickness ($\mu\text{m}$) | | % Neointima | | % Venous Muscularization | |
| --- | --- | --- | --- | --- | --- | --- |
|  | Adjusted p-value | Sig. | Adjusted p-value | Sig. | Adjusted p-value | Sig. |
| 8wk PBS vehicle vs. 8wk PBS imatinib | 0.741 | ns | 1.000 | ns | 1.000 | ns |
| 8wk PBS vehicle vs. 6wk HDM vehicle | $3.57 \times 10^{-3}$ | ** | $1.20 \times 10^{-4}$ | *** | $3.99 \times 10^{-4}$ | *** |
| 8wk PBS vehicle vs. 6wk HDM imatinib | $4.16 \times 10^{-4}$ | *** | $3.15 \times 10^{-4}$ | *** | $4.29 \times 10^{-3}$ | ** |
| 8wk PBS vehicle vs. 8wk HDM vehicle | $2.70 \times 10^{-5}$ | **** | $1.50 \times 10^{-5}$ | **** | $3.76 \times 10^{-5}$ | **** |
| 8wk PBS vehicle vs. 8wk HDM imatinib | $8.77 \times 10^{-5}$ | **** | $7.80 \times 10^{-6}$ | **** | $3.57 \times 10^{-4}$ | *** |
| 8wk PBS imatinib vs. 6wk HDM vehicle | $4.63 \times 10^{-4}$ | *** | $8.85 \times 10^{-4}$ | *** | $2.98 \times 10^{-4}$ | *** |
| 8wk PBS imatinib vs. 6wk HDM imatinib | $7.01 \times 10^{-5}$ | **** | $1.45 \times 10^{-3}$ | ** | $2.82 \times 10^{-3}$ | ** |
| 8wk PBS imatinib vs. 8wk HDM vehicle | $7.18 \times 10^{-6}$ | **** | $1.73 \times 10^{-4}$ | *** | $3.53 \times 10^{-5}$ | **** |
| 8wk PBS imatinib vs. 8wk HDM imatinib | $1.93 \times 10^{-5}$ | **** | $9.87 \times 10^{-5}$ | **** | $2.53 \times 10^{-4}$ | *** |
| 6wk HDM vehicle vs. 6wk HDM imatinib | 0.822 | ns | 1.000 | ns | 0.996 | ns |
| 6wk HDM vehicle vs. 8wk HDM vehicle | 0.626 | ns | 0.995 | ns | 0.951 | ns |
| 6wk HDM vehicle vs. 8wk HDM imatinib | 0.856 | ns | 0.974 | ns | 1.000 | ns |
| 6wk HDM imatinib vs. 8wk HDM vehicle | 1.000 | ns | 1.000 | ns | 0.797 | ns |
| 6wk HDM imatinib vs. 8wk HDM imatinib | 1.000 | ns | 0.998 | ns | 0.998 | ns |
| 8wk HDM vehicle vs. 8wk HDM imatinib | 0.998 | ns | 1.000 | ns | 0.915 | ns |

|  |  | Avg. Medial Thickness (μm) |  | % Neointima |  |
| --- | --- | --- | --- | --- | --- |
| Size Bin | Contrast (imatinib vs. vehicle) | Adjusted p-value | Sig. | Adjusted p-value | Sig. |
| 25–50 μm | 8wk PBS imatinib vs. 8wk PBS vehicle | 0.872 | ns | 1.000 | ns |
|  | 6wk HDM imatinib vs. 6wk HDM vehicle | 0.988 | ns | 0.733 | ns |
|  | 8wk HDM imatinib vs. 8wk HDM vehicle | 0.319 | ns | 0.602 | ns |
| 51–100 μm | 8wk PBS imatinib vs. 8wk PBS vehicle | 0.429 | ns | 1.000 | ns |
|  | 6wk HDM imatinib vs. 6wk HDM vehicle | 0.143 | ns | 0.979 | ns |
|  | 8wk HDM imatinib vs. 8wk HDM vehicle | 1.000 | ns | 1.000 | ns |
| 101–150 μm | 8wk PBS imatinib vs. 8wk PBS vehicle | 0.954 | ns | 1.000 | ns |
|  | 6wk HDM imatinib vs. 6wk HDM vehicle | 0.992 | ns | 0.881 | ns |
|  | 8wk HDM imatinib vs. 8wk HDM vehicle | 0.999 | ns | 0.997 | ns |

